# Proteomic and Transcriptomic Profiling Identifies Mediators of Anchorage-Independent Growth and Roles of Inhibitor of Differentiation Proteins in Invasive Lobular Breast Cancer

**DOI:** 10.1101/543132

**Authors:** Nilgun Tasdemir, Kai Ding, Kevin M. Levine, Tian Du, Emily A. Bossart, Adrian V. Lee, Nancy E. Davidson, Steffi Oesterreich

## Abstract

**BACKGROUND:** Invasive lobular carcinoma (ILC) is a histological subtype of breast cancer with distinct molecular and clinical features from the more common subtype invasive lobular carcinoma (IDC). We have previously shown that human ILC cells lines have a remarkably unique ability to grow in ultra-low attachment (ULA) suspension cultures as compared to IDC cells, the mediators of which remain unknown.

**METHODS:** Using flow cytometry and immunoblotting in human ILC and IDC cell lines, we measured levels of apoptosis and cell proliferation in attached (2D) and suspension (ULA) cultures. siRNA-mediated knockdown and pharmacological inhibitors were utilized to assess the effects of known regulators of anchorage-independence. Reverse Phase Protein Arrays and RNA-Sequencing were performed to identify novel proteomic and transcriptomic mediators of ULA growth in ILC cells.

**RESULTS:** We show that human ILC cell lines exhibit enhanced anoikis resistance and cell proliferation in ULA cultures as compared to IDC cells. Transient restoration of E-cadherin did not impact the 2D or ULA growth of human ILC cell lines, while transient E-cadherin knockdown in IDC cells partially rescued their growth defect in ULA culture. Inhibition of the Rho/ROCK, p120-catenin or YAP/Hippo pathways previously implicated in anoikis resistance did not have a major effect on the ULA growth of ILC cells. Proteomic comparison of ILC and IDC cell lines identified unique induction of PI3K/Akt and p90-RSK pathways in ULA culture in ILC cells. Transcriptional profiling uncovered unique upregulation of the Inhibitors of Differentiation family transcription factors *ID1* and *ID3* in ILC ULA culture, the knockdown of which diminished anchorage-independent growth. We find that *ID1* and *ID3* expression is higher in human ILC tumors as compared to IDC and correlated with a worse disease-specific survival uniquely in the ILC cohort.

**CONCLUSION:** Our comprehensive study of 2D and ULA growth in human ILC cell lines revealed anoikis resistance, cell proliferation and novel mediators of anchorage-independence and provides possible mechanistic insights and clinical implications for metastatic dissemination of ILC. High expression in human ILC tumors and association with clinical outcome implicate *ID1* and *ID3* as novel drivers and therapeutic targets for lobular breast cancer.

## Background

Invasive lobular carcinoma (ILC) is one of the major histological subtypes of breast cancer, which accounts for ∼10-15% of all cases [1]. Compared to the more common subtype invasive ductal carcinoma (IDC), ILC has a number of unique histological, molecular and clinical characteristics. ILC tumors exhibit single-file growth invading the surrounding stroma in a diffuse, linear pattern [2]. This unusual feature is largely attributed to the hallmark genetic loss of *CDH1*, which encodes the adherens junction protein E-cadherin [3-5]. Despite their favorable prognostic and predictive factors such as belonging mainly to the Luminal A (LumA) subtype and low proliferation [6], ILC tumors paradoxically exhibit more frequent long-term recurrences on endocrine therapy than IDC tumors [7, 8]. Furthermore, patients with ILC frequently present with metastatic dissemination to unusual anatomical sites such as the peritoneum, ovaries and gastrointestinal tract [1, 7], clinical features that are not currently well understood.

Most mammalian cells need to continually maintain contacts with extracellular matrix (ECM) to engage integrin receptors for sustained downstream signaling pathways such as Focal Adhesion Kinase (FAK) [9, 10]. In the absence of such cell-matrix interactions, cells undergo detachment-induced cell death, which is termed anoikis [11]. As part of their evolution, some cancer cells acquire an ability to survive in detached conditions, known as anoikis resistance [12]. This anchorage-independence ability allows cells that have escaped from the site of the primary tumor to survive in the blood stream and subsequently at foreign matrix environments at secondary organs [13, 14]. As such, anoikis resistance is believed to be an important contributor to tumor cell dissemination and a surrogate indicator of distant metastatic colonization ability [15, 16].

Anchorage-independence has previously been described in mouse models of lobular cancer, where combined somatic inactivation of p53 and E-cadherin induces metastatic ILC through induction of anoikis resistance [17, 18]. Furthermore, loss of other adherens junction proteins such as p120-catenin (p120) also induces metastatic progression of mouse ILC cells through ROCK1-mediated anoikis resistance [19-21]. Recently, loss of E-cadherin has additionally been linked to anoikis resistance through PI3K/Akt pathway activation in mouse ILC cells and the Estrogen Receptor (ER)-negative human ILC cell line IPH-926 [22]. However, anoikis resistance has not previously been investigated in ER-positive human ILC cell lines. Furthermore, these mouse studies focused only on anoikis resistance during anchorage-independent growth, without assessing the proliferation ability of cells growing in suspension.

We have recently published a comprehensive phenotypic characterization of human ER-positive ILC cell lines in 2D and 3D cultures, where we reported that ILC cells exhibit a remarkably unique ability to efficiently grow in ultra-low attachment (ULA) culture as compared to IDC cells [23]. Given the importance of anchorage-independence in tumor cell dissemination and metastasis [2, 13, 15, 24], herein we characterized the cellular and molecular mechanisms underlying the ability of human ILC cells to survive in detached conditions. Using a series of human IDC cell lines for comparison, our work revealed a combined mechanism of anoikis resistance and sustained cell proliferation driving the survival of human ILC cell lines in ULA culture. In addition to ruling out major roles for the previously described regulators of anoikis resistance, our molecular profiling studies uncovered novel mediators of ILC anchorage-independent growth that may constitute novel therapeutic targets for patients with lobular breast cancer.

## Methods

### Cell culture

MDA-MB-134-VI, MDA-MB-330, MCF-7, T47D, MDA-MB-231 and SKBR3 cells were obtained from the American Type Culture Collection. SUM44PE (SUM44) cells were purchased from Asterand and BCK4 cells were kindly provided by Britta Jacobsen, University of Colorado Anschutz, CO. Cell lines were maintained as previously described [23] in the following media (Life Technologies) with 10% FBS: MDA-MB-134 and MDA-MB-330 in 1:1 DMEM:L-15, MCF7 and MDA-MB-231 in DMEM, T47D in RPMI, BCK4 in MEM with non-essential amino acids (Life Technologies) and insulin (Sigma-Aldrich). SUM44 cells were maintained as described [25] in DMEM-F12 with 2% charcoal stripped serum and supplements. Cell lines were routinely tested to be mycoplasma free, authenticated by the University of Arizona Genetics Core by Short Tandem Repeat DNA profiling and kept in continuous culture for <6 months.

### Transient transfection, siRNAs, plasmids and drugs

ON-TARGETplus SMARTpool small interfering RNAs (siRNAs) were purchased from Dharmacon: ID1 (#L-005051-00-0005), ID3 (#L-009905-00-0005), ROCK1 (#L-003536-00-0005), p120 (#L-012572-00-0005), YAP (#L-012200-00-0005), non-targeting control (#D-001810-10-50). Cells were reverse-transfected with 1pmol/10nM of each siRNA in 96-well plates using Lipofectamine RNAiMAX (Life Technologies) and Opti-MEM (Thermo Fisher Scientific) following manufacturer’s instructions. pCDNA3 backbone was from Invitrogen and hE-cadherin-pcDNA3 was a gift from Barry Gumbiner (Addgene plasmid # 45769). Plasmids were forward transfected into cells using FuGENE 6 (Promega). BEZ-235, LY-294002, GSK-1120212, SCH-772984 and LJH-685 were purchased for Selleck Chemicals. Y-27632 was purchased from Sigma Aldrich.

### Cell viability and anchorage-independence assays

2D and ULA growth assays were performed as previously described [23]. ILC (15,000/96-well; 300,000/6-well) and IDC (5,000/96-well; 100,000/6-well) cells were seeded in regular (Thermo Fisher Scientific) or ULA (Corning Life Sciences) 96-well plates. Cells were assayed using CellTiter-Glo (Promega), Fluoreporter Blue Fluorometric dsDNA Quantitation Kit (Invitrogen) or PrestoBlue Cell Viability Reagent (Thermo Fisher Scientific). Data was captured on a Promega GloMax or Perkin Elmer plate reader.

### Cell Proliferation, Cell Cycle and Apoptosis Assays

Cells were seeded at 300,000/well in 6-well 2D and ULA plates in triplicates. For proliferation assays, cells were stained with 0.01μM carboxyfluorescein succinimidyl ester (CFSE; Thermo Fisher Scientific) for 20 minutes at room temperature in serum-free media on day 0. At the indicated time points, cells were harvested by trypsinization of 2D cultures in plates and ULA cultures in tubes for 5 minutes at 37°C, followed by neutralization with serum-containing media and washing with PBS. Ki67/7-AAD staining was performed using the FITC Mouse Anti-Ki67 set (#556026; BD Biosciences) following manufacturer recommendations and gating on cells stained with an isotype control antibody. For cell cycle analysis, cells were stained with Hoechst (Thermo Fisher Scientific) at 20mg/ml for 30 minutes at 37°C and then briefly with Propidium Iodide (BD Biosciences) to gate on viable cells. For apoptosis assays, cells were stained with APC-Annexin V (BD Biosciences; #550474) and PI in 1X Annexin binding buffer for 15 minutes at room temperature. Samples were acquired on an LSR II Flow cytometer (BD Biosciences) and analyzed using BD FACSDiva software (BD Biosciences).

### Immunoblotting and Reverse Phase Protein Arrays (RPPA)

Immunoblots were performed as previously described [23] using 5% milk powder for blocking and developed using ECL (Sigma-Aldrich). Details of the antibodies used in **Additional file 1: Table S1**. RPPA was performed as previously described [25]. Samples were collected in MD Anderson RPPA lysis buffer and assessed at the Functional Proteomics Core of MD Anderson. Rawlog2 RPPA data is included in **Additional file 2: Table S2.** Normalized log2 median-centered values were used to generate heat maps.

### RNA extraction, quantitative PCR and RNA-Sequencing (RNA-Seq)

RNA extraction and qRT-PCRs were done as previously described [23]. RNA-Seq was performed as previously described [26] using Illumina HiSeq 2000. Raw sequence data were mapped to hg38 genome (ensemble release version 82) and gene counts were quantified with Salmon (version 0.6.0) [27] using default settings. Differentially expressed (DE) analysis was performed with R package DESeq2 [28] using the following criteria: absolute log2(fold change)>log2(1.5) and Benjamini-Hochberg adjusted p-value <0.05. The complete list of DE genes is available in **Additional files 3-6: Table S3-S6**. Venn diagrams were generated using the online Intervene tool [29].

### Survival analyses

Survival analyses was performed using the METABRIC dataset [30] as previously described [31], using data downloaded from the Synapse software platform (syn1688369; Sage Bionetworks, Seattle, WA, USA).

### Statistical analysis

Data analysis was performed using GraphPad Prism. Data is presented as mean +/− standard deviation or standard error of means as indicated. Statistical tests used for each figure are indicated in the respective figure legends.

## Results

### Anoikis resistance and cell proliferation during ILC anchorage-independent growth

We have previously shown that the ER-positive human ILC cell lines MDA-MB-134 (MM134), SUM44PE (SUM44), MDA-MB-330 (MM330) and BCK4 can grow efficiently in ULA suspension culture, as compared to the limited growth of the ER+ human IDC cell lines MCF7 and T47D, and the ER-negative human IDC cell line MDA-MB-231 (MM231) under the same conditions [23]. As these results had been obtained using the CellTiter-Glo luminescent cell viability reagent, here we repeated these experiments with the FluoReporter fluorescent dsDNA assay, the output of which does not rely on the metabolic activity of cells. Consistent with our previous results, we observed a superior growth ability of ILC versus IDC cell lines in ULA culture and additionally showed that the ER-negative, HER2-positive human IDC cell line SKBR3 [32] also exhibits limited ULA growth (**Additional file 7: Figure S1A-C**).

Having confirmed the differential growth of ILC and IDC cells in ULA culture, we next assessed the levels of anoikis in these cell types. Annexin V and propidium iodide (PI) flow cytometry (FACS) analysis of MM134, SUM44, MCF7 and T47D cells grown in 2D or ULA showed that both ILC and IDC cell lines display high levels of anoikis resistance (**Figure 1A, B**). Quantification of the double-negative, viable cells indicated no anoikis in MM134 and SUM44 and only ∼20% (MCF7) or ∼5% (T47D) in the IDC cells (**Figure 1C, D**). This anoikis phenotype was further assessed by immunoblotting, which revealed an increase in cleaved PARP (lower band) in ULA versus 2D only in MCF7 cells (**Figure 1E)**. Furthermore, we confirmed these findings in additional cell lines and observed generally lower levels of anoikis in ILC versus IDC cells (**Additional file 7: Figure S2**).

**Fig. 1.**
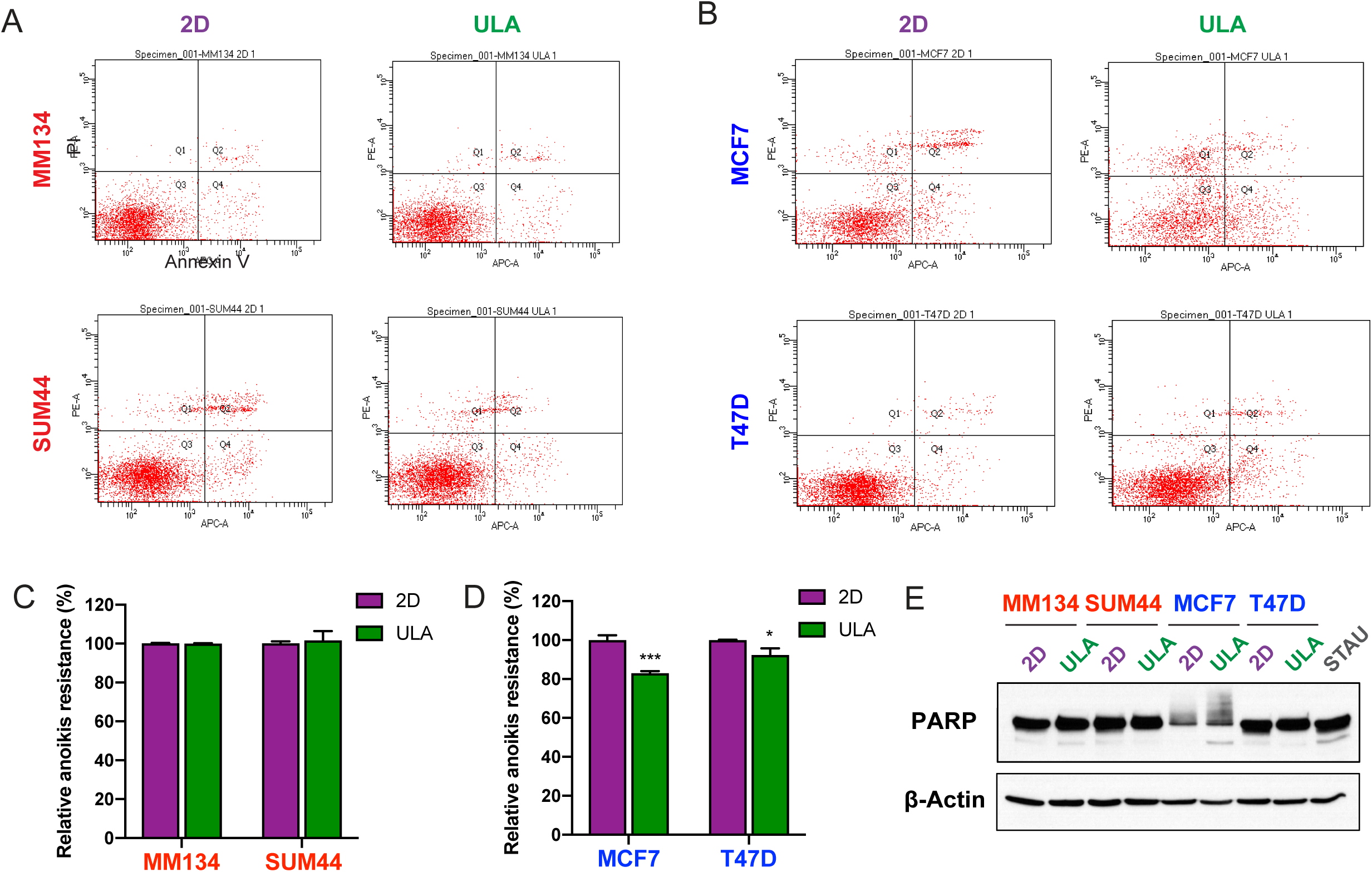
Anoikis resistance of human ILC and IDC cell lines. **a-b**. Representative Annexin V and PI FACS staining plots of the (**a**) ILC (red) cell lines MM134 (top) and SUM44 (bottom) and (**b**) IDC (blue) cell lines MCF7 (top) and T47D (bottom) after 4 days in 2D (left; purple) or ULA (right; green) culture. **c-d** Quantification of the anoikis resistant (Annexin V-/PI-) population in (**c**) ILC and (**d**) IDC cell lines. Data is displayed as mean percentage +/− standard deviation relative to the 2D condition in each cell line. Graphs show representative data from two experiments (n=3). p-values are from t-tests. * p ≤ 0.05; *** p ≤ 0.001. **e** Immunoblotting for PARP in ILC and IDC cell lines after 2 days in 2D or ULA culture. STAU: positive control from T47D cells treated with 1 μM Staurosporine for 5 hours. β-Actin was used as a loading control

Given the large differences in the viabilities of ILC and IDC cells in ULA conditions (see **Figure S1**) despite generally high levels of anoikis resistance in both cell types (see **Figure 1** and **Additional file 7: Figure S2**), we reasoned that they might exhibit different levels of proliferation in ULA conditions. FACS-based Hoechst staining revealed similar cell cycle profiles for MM134 and SUM44 in 2D and ULA, whereas MCF7 and T47D exhibited more cells arrested in G0/G1, concomitant with a decrease in the percentage of cells in the S and G2/M phases in ULA compared to 2D conditions (**Figure 2A-D**). We confirmed these findings by additional FACS analyses, which showed more CFSE-retaining IDC cells in ULA (**Figure 2E, F**), as well as lower Ki67 positivity in these cells as compared to 2D (**Figure S3**), despite similar levels for ILC cells in both conditions and assays. Collectively, these data indicate that the superior viability of human ILC cells in ULA conditions compared to IDC cells is due to a combined mechanism of anoikis resistance and sustained cell proliferation.

**Fig. 2.**
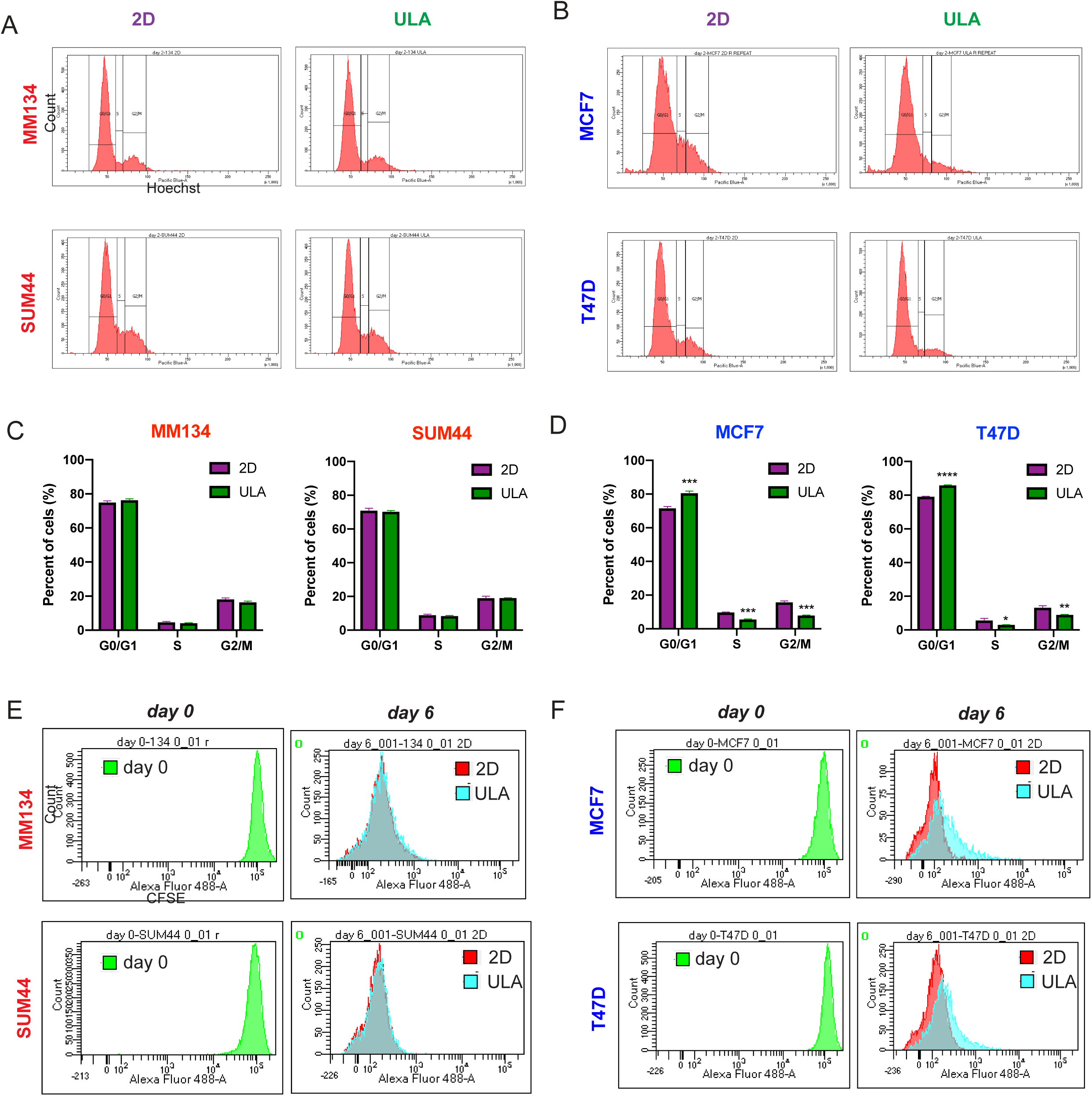
Cell cycle and cell proliferation in ILC and IDC cell lines in 2D and ULA culture. **a-b** Representative FACS plots from Hoechst staining of the (**a**) ILC (red) cell lines MM134 (top) and SUM44 (bottom) and (**b**) IDC (blue) cell lines MCF7 (top) and T47D (bottom) after 2 days in 2D (left; purple) or ULA (right; green) culture. **c-d** Quantification of the cells in the indicated phases of the cell cycle based on the gating in (**a, b**) in (**c**) ILC and (**d**) IDC cell lines. Data is displayed as mean percentage +/− standard deviation (n=3). p-values are from t-tests. * p ≤ 0.05; ** p ≤ 0.01; *** p ≤ 0.001; **** p ≤ 0.0001. **e-f** CFSE FACS plots of the (**e**) ILC cell lines MM134 (top) and SUM44 (bottom) and (**f**) IDC cell lines MCF7 (top) and T47D (bottom) after initial labeling (day 0; left) and 6 days (right) in 2D or ULA culture shown as overlays

### Roles of known regulators of anchorage-independence in ILC ULA growth

Anoikis resistance has previously been attributed to the absence of adherens junctions, which is due to the hallmark genetic loss of E-cadherin (encoded by the *CDH1* gene) in mouse ILC cells [17, 18]. To test the involvement of E-cadherin loss in the anchorage-independent growth of human ILC cell lines, we transiently overexpressed *CDH1* in MM134 and SUM44. Re-introduction of E-cadherin did not affect the growth of these ILC cell lines in either 2D or ULA culture (**Figure 3A-C**). As a complementary approach, we also transiently knocked down *CDH1* in MCF7 and T47D cells using siRNAs. The diminished levels of E-cadherin in these IDC cells led to a rounded cell morphology and partially rescued the growth in ULA culture, but not fully to the levels of growth in 2D culture (**Figure 3D-F**). These data are also consistent with our observation of limited ULA growth in the E-cadherin defective cell lines MM231 (*CDH1* promoter hypermethylation) and SKBR3 (homozygous *CDH1* deletion) [33].

**Fig. 3.**
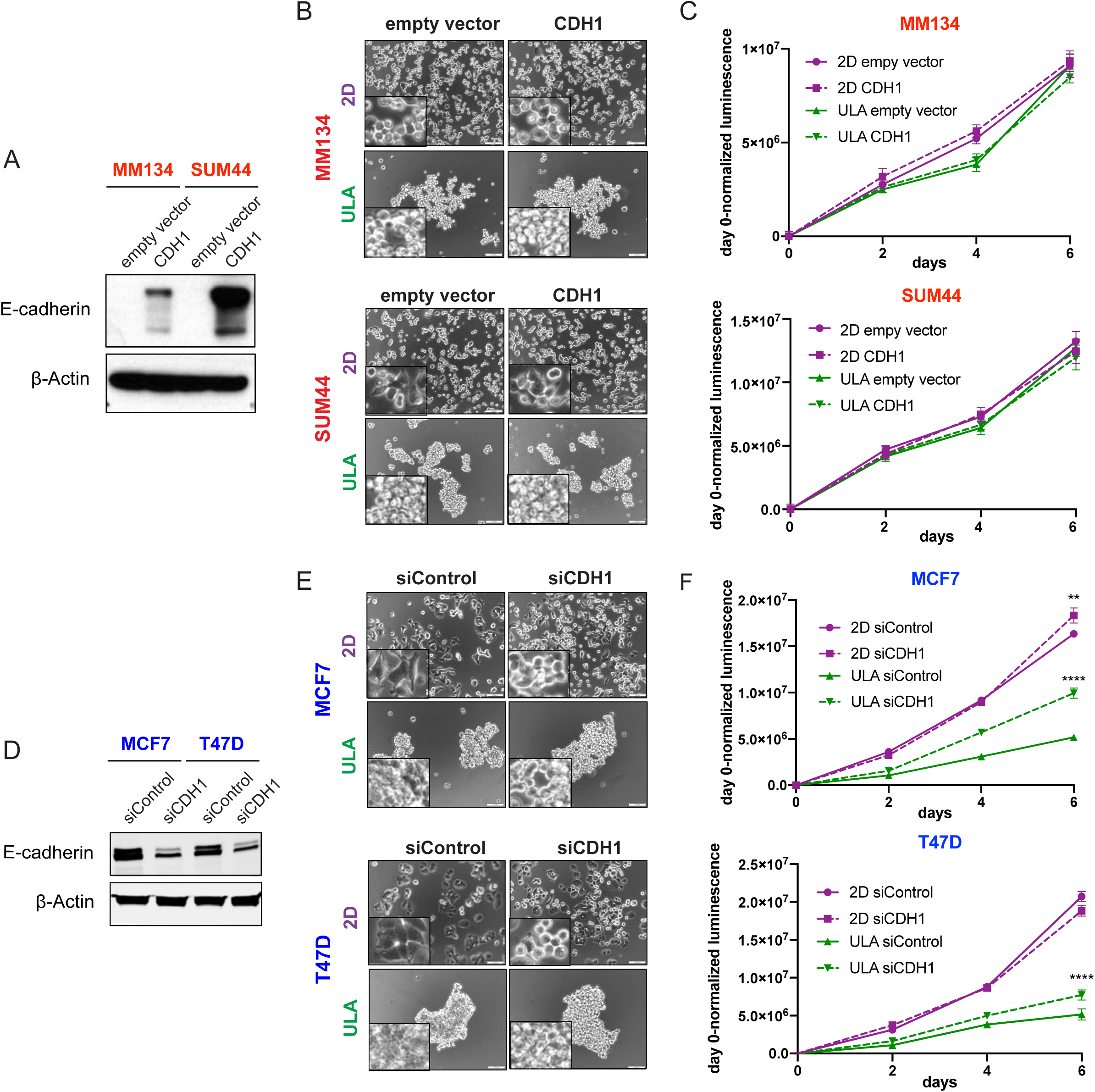
Effects of CDH1 restoration in ILC and CDH1 knockdown in IDC cell lines on cell morphology and viability in 2D and ULA culture. **a-f** Immunoblotting for E-cadherin (**a, d**), morphology (**b, e**) and growth (**c, f**) in 2D (purple) or ULA (green) culture in the (**a, c**) ILC (red) cell lines MM134 (left; top) and SUM44 (right; bottom) transiently transfected with an empty or CDH1 overexpression vector and (**d-f**) IDC (blue) cell lines MCF7 (left; top) and T47D (right; bottom) transiently transfected with a control or CDH1 siRNA. β-Actin was used as a loading control. Graphs show representative data from two experiments (n=6). p-values are from two-way ANOVA comparison of (**c**) empty vector and CDH1 or (**f**) siControl and siCDH1 in 2D and ULA culture separately. ** p ≤ 0.01; **** p ≤ 0.0001

Besides E-cadherin, a number of other genes and pathways such as Rho/ROCK, p120 and YAP/Hippo have also been previously implicated in anoikis resistance [19-21, 34-36]. Thus, we next tested the effect of the ROCK inhibitor Y-27632 on the anchorage independent growth of ILC and IDC cell lines and generally observed very similar dose response curves in 2D and ULA cultures (**Figure 4A, B** and **Additional file 7: Figure S4A, B**). Of note, a specific differential effect in ULA versus 2D was observed in MCF7 (**Figure 4B, C**) and MM330 (**Additional file 7: Figure S4A, B**) cells only at 10 μM, a concentration almost universally used in literature [19, 37, 38]. Consistent with previously reported effects of ROCK inhibition [39-41], Y-27632 at this dose led to a generally elongated morphology in all cell lines in 2D culture but to significantly tighter colony formation in only MCF7 (**Figure 4D**) and MM330 (**Additional file 7: Figure S4C**) cells in ULA. In addition, we also used siRNAs to transiently knockdown ROCK1, p120 and YAP in MM134 and SUM44 cells, which generally resulted in little to no effect on growth in 2D or ULA culture (**Additional file 7: Figure S5**). The most substantial reduction in viability was observed with siYAP in MM134 cells in ULA, which was stronger than in 2D; however, this finding did not translate to SUM44 cells. Combined, these data suggest that the regulation of anchorage-independent growth in human ILC cell lines differs from that of human IDC and mouse ILC cells.

**Fig. 4.**
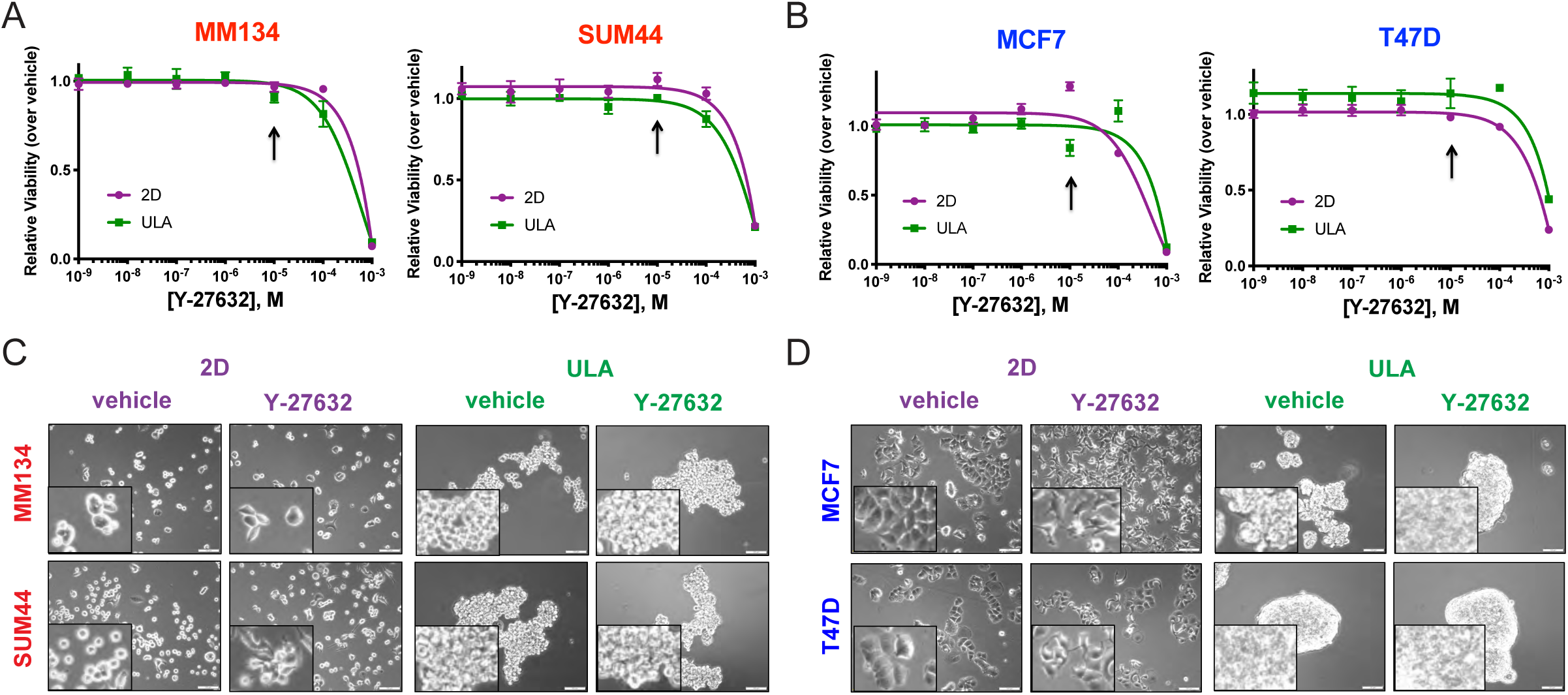
Effects of ROCK inhibition on the viability and morphology of ILC and IDC cell lines in 2D and ULA culture. **a-b** Dose response curves of the (**a**) ILC (red) cell lines MM134 (left) and SUM44 (right) and (**b**) IDC (blue) cell lines MCF7 (left) and T47D (right) treated with the indicated doses of the Y-27632 ROCK inhibitor in 2D (purple) or ULA (green) culture after 4 days. Arrows indicate the dose used for the morphology pictures in (**c-d**). **c-d** Morphologies of the (**c**) ILC and (**d**) IDC cell lines from (**a-b**) treated with vehicle or 10 μM (10^−5^ M) Y-27632 for 4 days in 2D or ULA culture. Insets show higher magnification images. Scale bar: 100 μM

### Proteomic mediators of ILC anchorage-independent growth

Having ruled out a substantial role for E-cadherin, ROCK, p120 and YAP in the anchorage independent growth of human ILC cell lines via our candidate approach, we next embarked on an unbiased proteomic analysis using Reverse Phase Protein Arrays (RPPA) using extracts from ILC and IDC cell lines after a 24-hour ULA culture (**Figure 5A, Additional file 2: Tables S2** and **Additional file 7: Figure S6**). As also confirmed by immunoblotting (**Figure 5B**), we observed reduced FAK phosphorylation in both ILC and IDC cell lines after a 24-hour ULA culture consistent with inactive integrin signaling in the absence of matrix. This RPPA analysis also revealed sustained PI3K/Akt pathway activation in SUM44 cells, which was downregulated in IDC cells in ULA (**Figure 5A, B**). Interestingly, while the PDK1-induced phosphorylation of Akt (T308) and the Akt target PRAS40 (T246) were sustained in ULA culture in MM134 cells, the mTOR-induced phosphorylation of Akt (S473) and downstream PI3K/Akt targets were paradoxically downregulated. Given the longer duration of our growth assays, we also assayed the same pathways after 4 days of 2D and ULA culture and observed upregulation in MM134 and downregulation in SUM44 cells in ULA at this time point, suggesting that they were highly dynamically regulated (**Additional file 7: Figure S7A**), consistent with the complexity of the PI3K/Akt pathway [42].

**Fig. 5.**
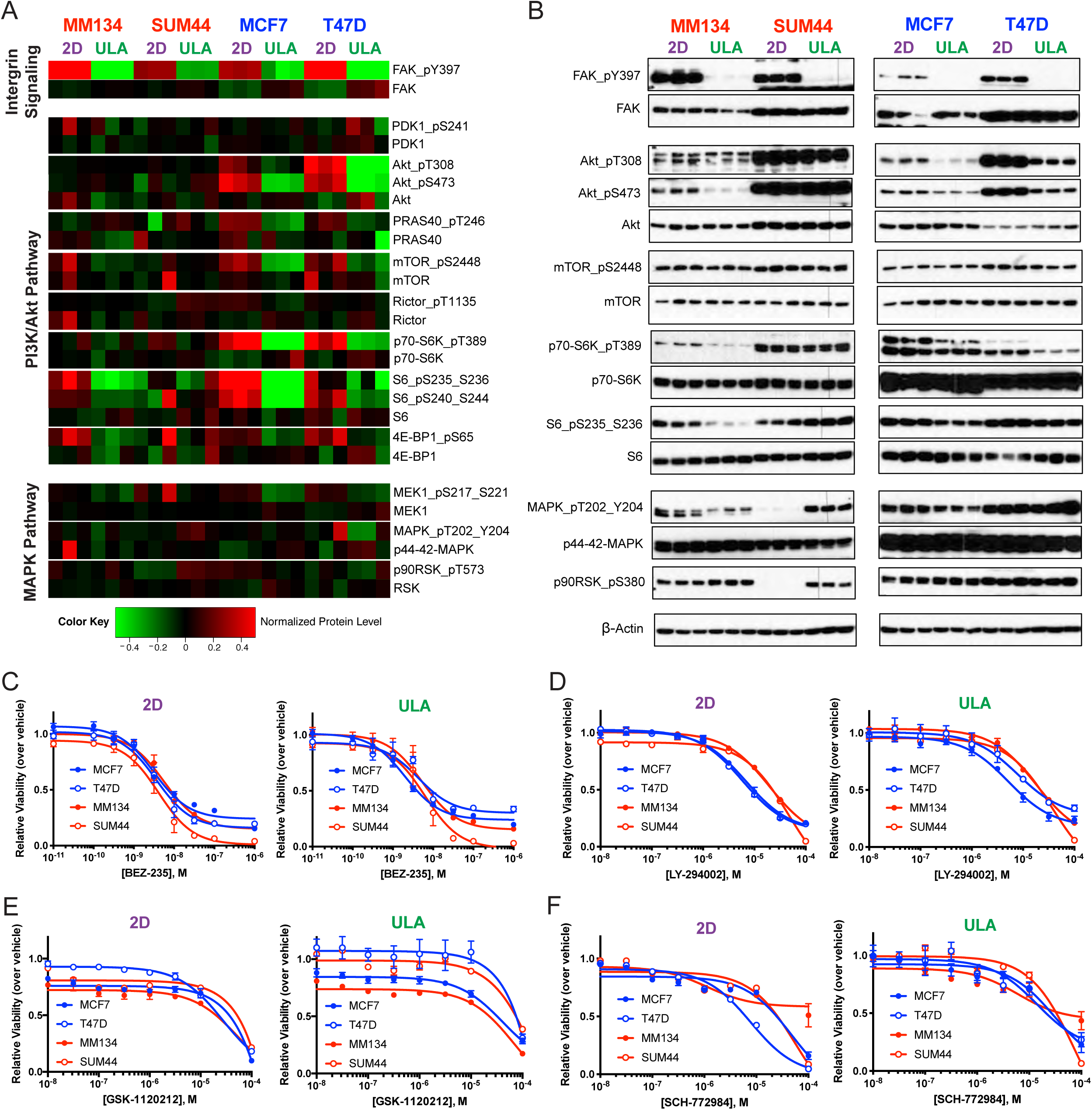
Proteomic profiling and drug treatments of ILC and IDC cell lines in 2D and ULA culture. **a-b** RPPA (**a**) and Western blot (**b**) analyses of the ILC (red) cell lines MM134 and SUM44 and IDC (blue) cell lines MCF7 and T47D grown in 2D (purple) or ULA (green) culture for 24 hours for the indicated pathways and proteins. Three biological replicates are displayed for each condition. β-Actin was used as a loading control. **c-f** Dose response curves of the ILC and IDC cell lines from (**a**) treated with the indicated doses of the (**c**) PIK3CA/mTOR inhibitor BEZ-235, (**d**) PIK3CA inhibitor LY-294002, (**e**) MEK inhibitor GSK-1120212 and (**f**) MAPK inhibitor SCH-772984 in 2D (purple; left) or ULA (green; right) culture after 4 days.

Another pathway of interest revealed by our RPPA profiling was Ras/MAPK, as we observed increased phosphorylation of the downstream effector p90RSK in ILC ULA culture, which was not observed in IDC cells (**Figure 5A, B**). Interestingly, this phosphorylation was independent of MEK and MAPK activation, as increased p90RSK phosphorylation was observed in MM134 cells in ULA despite decreased upstream signaling. Similarly, T47D cells did not exhibit induction of p90RSK phosphorylation in ULA in the presence of concomitant upstream MAPK signaling. These results suggested an uncoupling of Ras/MAPK activity and downstream p90RSK phosphorylation in ULA conditions.

Next we tested the effects of pharmacological inhibitors targeting PI3K/Akt and Ras/MAPK pathways and did not observe significant shifts in the overall dose response curves between the different cell lines and culture conditions. Of note, the ILC cell lines showed the strongest sensitivity to the PI3K/mTOR dual inhibitor BEZ-235 in ULA at the highest doses; however, similar effects were also observed in 2D (**Figure 5C**). Interestingly, LY-294002, which only targets PI3K, had the strongest efficacy in the IDC cell lines in both conditions (**Figure 5D**), in agreement with the previously reported sensitizing effects of their activating PI3K mutations [22, 43-45]. Treatment with the MEK inhibitor GSK-1120212 showed the most efficacy in MM134 cells in both 2D and ULA (**Figure 5E**), consistent with the inactivating *MAP2K4* mutation in this cell line [46]. Conversely, targeting the same pathway further downstream with the MAPK inhibitor SCH-772984 had the least efficacy in this cell line (**Figure 5F**), supporting the highly complex and uncoupled regulation of this pathway. Finally, treatment of ILC and IDC cell lines with the p90RSK inhibitor LJH-685 also did not reveal a significant difference between the dose response curves, although full inhibition could not be achieved even at the highest doses (**Additional file 7: Figure S7B**).

### Transcriptomic mediators of ILC anchorage-independent growth

Given the lack of effective and specific blockage of the ILC ULA growth by the tested pharmacological inhibitors, we next performed RNA-Seq to investigate the transcriptional outputs of different culture conditions. To focus on acute transcriptional changes and to mirror the proteomic profiling experiments, we analyzed cells grown in 2D or ULA culture for 24 hours. Despite highly similar overall transcriptional profiles as revealed by Principal Component Analysis (PCA) (**Figure 6A**), RNA-Seq identified a number of differentially regulated protein-coding and long non-coding genes in the two culture conditions (**Figure 6B, Additional file 7: Figure S8A** and **Additional files 3-6: Tables S3-6**). We focused on the ULA-upregulated transcripts in both MM134 and SUM44 cells, which were not shared with MCF7 or T47D, including a total of 10 genes (**Figure 6B**).

**Fig. 6.**
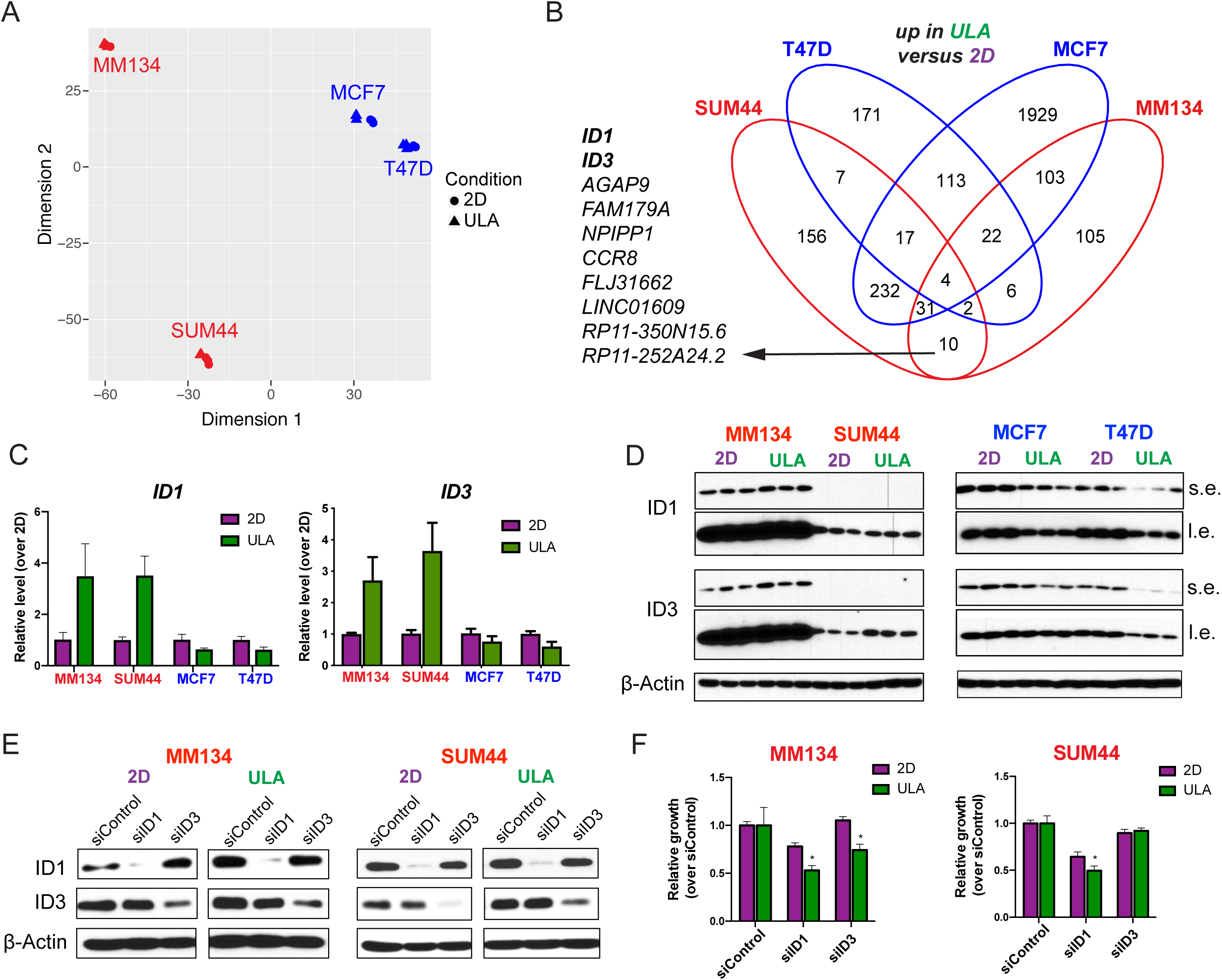
Transcriptomic profiling of ILC and IDC cell lines in 2D and ULA culture. **a** PCA of transcriptomic data from the ILC (red) and IDC (blue) cell lines grown in 2D (circle) or ULA (triangle) culture for 24 hours using the top 5000 most variable genes ranked by interquartile range. **b** Venn diagrams showing the overlap between the genes upregulated in ULA (green) culture as compared to 2D (purple) in the cells from (**a**). The list on the left shows the 10 genes commonly upregulated in the two ILC but not the IDC cell lines. **c-d** qRT-PCR (**c**) and immunoblotting (**d**) validation of the ID1 and ID3 upregulation in ULA culture in the ILC (left) but not IDC (right) cell lines. Data is displayed as mean +/− standard error relative to the 2D condition in each cell line. Graphs show data from three biological replicates. β-Actin was used as a loading control. s.e: short exposure. l.e: long exposure. **e-f** ID1 and ID3 immunoblotting (**e**) and growth (**f**) of MM134 (left) and SUM44 (right) cell lines 4 days after transient transfection with the indicated siRNAs. Data is displayed as mean +/− standard deviation relative to siControl in each condition in each cell line (n=6). p-values are from t-tests between 2D and ULA for each siRNA. * p ≤ 0.05

Given that *ID1* and *ID3* both encode transcription factors from the inhibitors of differentiation family of proteins, which have previously been implicated in metastasis [47, 48], we decided to follow up on these genes. We initially validated the ULA-induction of ID1 and ID3 at both the transcript (**Figure 6C**) and protein (**Figure 6D**) levels in ILC cell lines, which additionally revealed reciprocal downregulation in the IDC cells in ULA. Next we used siRNAs to knockdown *ID1* and *ID3* in MM134 and SUM44 cells (**Figure 6E** and **Additional file 7: Figure S8B, C**). Transient inhibition of both *ID1* resulted in reduced cell growth in both MM134 and SUM44 cells, with stronger effects in ULA versus 2D culture, while *ID3* knockdown resulted in a similar phenotype only in MM134 cells (**Figure 6F).** These data implicate *ID1* and *ID3* as novel drivers of ILC anchorage-independent growth and potential therapeutic targets.

### *ID1* and *ID3* expression and correlation with clinical outcome in human breast tumors

To assess the clinical relevance of our findings, we analyzed the RNA-Seq data from the human breast tumors in The Cancer Genome Atlas (TCGA) collection [6], which revealed higher *ID1* and *ID3* mRNA expression in both ER-positive and LumA ILC versus IDC tumors (**Figure 7A**). Similar results were observed in both ER-positive and LumA tumors from the METABRIC cohort [30] (**Figure 7B**). Finally, high combined *ID1* and *ID3* expression is correlated with significantly lower disease-specific survival in the LumA ILC but not LumA IDC patients in the METABRIC cohort (**Figure 7C**). Combined, these data suggest that *ID1* and *ID3* might be involved in the disease progression of patients with ILC, which warrants further investigation.

**Fig. 7.**
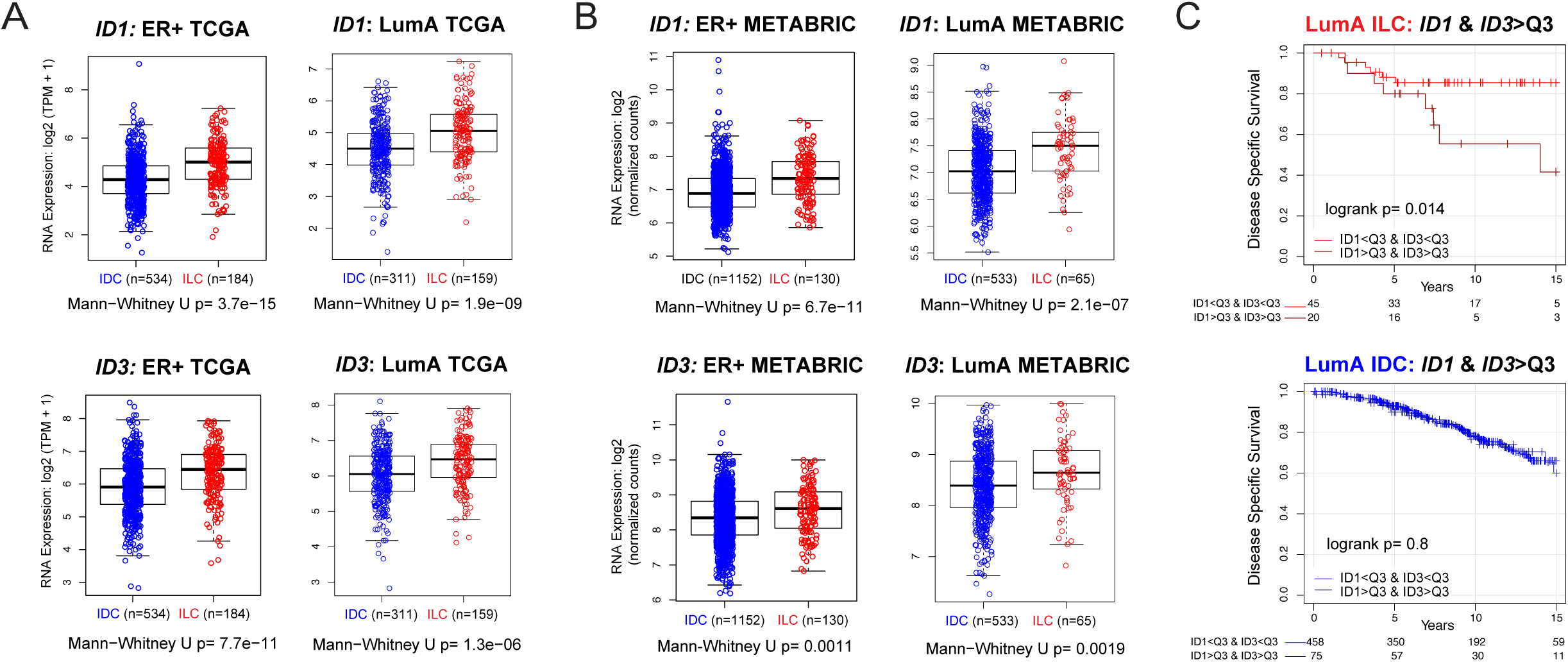
*ID1* and *ID3* expression in human breast tumors and correlation with survival. **a-b** mRNA levels of *ID1* (top) and *ID3* (bottom) in ER-positive (left) and LumA (right) ILC (red) and IDC (blue) tumors from the (**a**) TCGA and (**b**) METABRIC cohorts. p values are from Mann-Whitney U test. **c** Disease-specific survival curves for combined ID1 and ID3 expression in Luminal A ILC (top) and IDC (bottom) patients from the METABRIC cohort divided by third quartile (Q3) levels. p-values are from log-rank test

## Conclusion

ILC is a unique histological subtype of breast cancer that exhibits distinct molecular and clinical features from IDC [1]. Our previous work has identified a unique anchorage-independence ability of human ILC cell lines in ULA suspension culture [23]. Herein, we further characterized this interesting phenotype and uncovered a unique, combined mechanism of anoikis resistance and sustained cell proliferation, along with novel mediators of growth in detached culture, which were not shared with IDC cells. While anoikis resistance has been the major focus of existing work on anchorage-independence [11, 15, 16], the contribution of cell cycle progression and cell proliferation to this phenotype has been much less studied [12, 49-51]. Importantly, anoikis resistance and sustained cell proliferation during detached growth had not previously been studied in human ER-positive ILC cells.

Given the established role of anchorage-independence in tumor cell dissemination [2, 13, 15, 24], our findings may have a number of important clinical implications. Although ILC and IDC tumors both exhibit metastases, ILC tumors are associated with more frequent late recurrences. From this perspective, the superior ability of ILC cells to survive in detached conditions might allow them to persist at low levels of proliferation in foreign matrix conditions for extended periods of time. Furthermore, another unique feature of ILC tumors is their colonization of unique anatomical sites. It will be important to further study the matrix compositions of different metastatic organs and assess whether the ILC-specific sites are less permissive for re-establishing cell-ECM contacts, which would favor the survival of detached ILC versus IDC cells given our *in vitro* data and provide insights into metastatic organotropism.

Our findings indicate that anchorage-independence of human ILC cell lines differs from that of human IDC and mouse ILC cells in several ways. In contrast to their previously reported roles in anoikis resistance of mouse ILC cells [17-21], inhibition of E-cadherin and ROCK did not have major effects in human ILC ULA growth. Interestingly, while transient *CDH1* knockdown in IDC cells resulted in a rounded morphology, re-introduction of E-cadherin into ILC cells did not reconstitute the adherens junctions, likely due to the heterogeneous pool of transfected cells. Although our efforts of stable *CDH1* overexpression in human ILC cell lines have proven challenging so far, revisiting the role of E-cadherin in anchorage-independence in homogeneous cell populations with restored adherens junctions will be important. Nevertheless, knockdown of p120, the effector molecule downstream of E-cadherin loss previously implicated in mouse ILC anoikis resistance [19-21, 34], also did not significantly impact human ILC ULA growth. Similarly, inhibition of YAP, which has been linked to anoikis resistance and metastasis [34-36], had a minor differential effect in ULA only in MM134 cells, suggesting the involvement of additional, unique mediators.

As we previously reported [23], despite their efficient growth in ULA suspension culture, human ILC cell lines form very loose and slow growing colonies in 3D ECM gels or soft agar. Interestingly, a recent study using human IDC cell lines reported a strong correlation between the ability to grow in ULA and to form colonies in soft agar [52]. These data further highlight the differences in anchorage-independence between ILC and IDC and support the existence of unique mechanisms in the former. Although we have mainly focused on anoikis resistance and sustained cell proliferation in ILC ULA growth, the levels of detachment-induced death and cell cycle arrest were rather low in the IDC cells and did not fully account for the large ULA growth differences observed between the two subtypes. Thus, additional forms of non-apoptotic cell death such as entosis, cornification, and necrosis, as well as cell survival mechanisms such as autophagy and metabolic reprogramming, all of which have previously been linked to anchorage-independence [53-57], might be involved as parallel mechanisms and warrant further investigation.

Our molecular profiling experiments revealed few ULA-induced genes and pathways that were common to both ILC cell lines and not shared with IDC cells. The heterogeneity in the ULA-triggered transcriptional and proteomic changes, as well as in the timing of signaling activation, between MM134 and SUM44 cells suggests that these two cell types might potentially represent different ILC subtypes. This hypothesis is further supported by their highly separated clustering in our PCA analysis and harboring of different mutations [23, 46, 58], which might allow convergent but non-overlapping mechanisms of adaptation to detached growth. Although the 24-hour time point of suspension culture had previously been successfully utilized to uncover transcripts induced by detachment in triple negative breast cancer [59], the highly dynamic signaling we observed in ILC justifies further profiling studies focusing on earlier and later time points. In addition, as the RPPA only covers ∼220 total and phospho-proteins, more comprehensive assays such as Mass Spectrometry coupled with a complex systems biology approach are more likely to help fully understand the multi-dimensional regulation of signaling pathways unique to ILC suspension culture.

Although we detected ILC-unique ULA-induced PI3K/Akt activation, we did not observe significant shifts in the dose response curves to a PI3K/mTOR dual inhibitor. These data are in agreement with a recent report showing similar dose responses in adherent and suspension culture to Akt inhibitors in ER-negative mouse and human ILC cell lines [22]. While we noted stronger effects of the PI3K/mTOR dual inhibitor in ILC versus IDC cells in ULA only at the highest doses, a single PI3K inhibitor was conversely more effective in IDC versus ILC cell lines in both culture conditions despite decreased PI3K/Akt signaling in ULA, likely due to the PI3K mutations in MCF7 and T47D [22, 43-45]. Furthermore, we observed ILC-unique ULA-induced phosphorylation of p90RSK; however, a p90RSK inhibitor did not reveal any significantly differential sensitivity between the cell types and culture conditions. Given that full inhibition could not be achieved even at the highest doses of this compound, this finding warrants further investigation with more potent inhibitors. Interestingly, p90RSK has previously been implicated in soft agar and Matrigel growth of MM134 and mouse ILC cell lines downstream of FGFR1 and MEK/MAPK [60]. Here we report that ULA culture induces ILC-unique p90RSK phosphorylation, which is uncoupled from upstream MEK/MAPK signaling in the *MAP2K4* and *K-RAS* mutant MM134 cells [46, 58].

Our RNA-Seq profiling identified ILC-unique ULA-induction of transcripts encoding the inhibitor of differentiation family of proteins ID1 and ID3, which we found to be reciprocally downregulated in IDC ULA culture. *ID1* and *ID3* have previously been characterized as part of a common murine and human lung metastatic signature in triple negative breast cancer cells [48, 61] and extensively validated as regulators of metastasis [48, 62, 63]; however, they have not previously been studied in ILC. In our functional studies, siRNA-mediated knockdown of *ID1* and *ID3* in ILC cells resulted in reduced viability, which was stronger in ULA versus 2D. This dual effect in both culture conditions suggests that ID1 and ID3 inhibition may be effective on both the attached growth in primary tumors, as well as on the detached growth of disseminated cancer cells. Our findings are consistent with the previously reported roles of ID1 and ID3 in sustaining the proliferation of tumor cells during lung metastatic colonization, likely due to their negative regulation of basic-helix-loop-helix (bHLH) transcription factors [48].

Finally, we discovered significantly higher *ID1* and *ID3* transcript levels in human ILC tumors from both the TCGA and the METABRIC collections, which was consistent in both the ER-positive and the LumA subsets. Furthermore, combined expression of *ID1* and *ID3* is correlated with significantly worse disease-specific survival in the ILC but not IDC LumA METABRIC cohorts, suggesting a unique role in ILC. Collectively, these data implicate ID1 and ID3 as novel drivers and therapeutic targets in disease progression in ILC. Given the difficulty of blocking protein-protein interactions with bHLH transcription factors, most currently available inhibitors targeting ID1 and ID3, such as the ID1-degrader C527 [64] and Cannabidiol [65], are highly non-specific. More precise targeting strategies such as recently developed peptide-conjugated antisense oligonucleotides [66] should allow combined treatment with endocrine therapies to ultimately improve the clinical outcomes of patients with invasive lobular breast cancer.

## Declarations

### Ethics approval and consent to participate

Not applicable.

### Consent for publication

Not applicable.

### Competing interests

The authors have nothing to disclose.

## Funding

The work is in part funded by a Department of Defense Breakthrough Fellowship Award to NT (BC160764), Shear Family Foundation grant and Susan G. Komen Leadership grants to SO (SAC160073) and AVL (SAC110021), and Breast Cancer Research Foundation grants to NED and SO. KML was supported by an individual fellowship from the NIH/NCI (5F30CA203095). TD was supported by a China Scholarship Council award through Tsinghua School of Medicine, Beijing, China. EAB was supported by a National Institutes of Health (NIH) Ruth L. Kirschstein Award (1F31CA203055-01).

## Authors’ Contributions

***Conception and design:*** N. Tasdemir, NE. Davidson, S. Oesterreich

***Development of methodology:*** N. Tasdemir, AV. Lee, S. Oesterreich

***Acquisition of data (performed experiments, processed data, etc.):*** N. Tasdemir, K. Ding, EA. Bossart

***Analysis and interpretation of data (e.g. biological interpretation, statistical analysis, computational analysis):*** N. Tasdemir, K. Ding, KM. Levine, T. Du, AV. Lee, NE. Davidson, S. Oesterreich

***Writing, review and/or revision of the manuscript:*** N. Tasdemir, K. Ding, KM. Levine, T. Du, EA. Bossart, AV. Lee, NE. Davidson, S. Oesterreich

***Study supervision:*** AV. Lee, NE. Davidson and S. Oesterreich

## Acknowledgments

The authors thank Dr. Jennifer Atkinson for constructive comments during data interpretation and Jian Chen for outstanding technical support.

