## Supplemental Figures S1-8 and figure legends for "Proteomic and Transcriptomic Profiling Identifies Mediators of Anchorage-Independent Growth and Roles of Inhibitor of Differentiation Proteins in Invasive Lobular Breast Cancer"

**Supplementary Table and Figure Legends**

**Additional file 1: Table S1.** List of antibodies and primers used

**Additional file 2: Table S2.** Raw log2 RPPA data

**Additional file 3: Table S3.** Differentially regulated genes in ULA versus 2D in MM134 cells

**Additional file 4: Table S4**. Differentially regulated genes in ULA versus 2D in SUM44 cells

**Additional file 5: Table S5.** Differentially regulated genes in ULA versus 2D in MCF7 cells

**Additional file 6: Table S6.** Differentially regulated genes in ULA versus 2D in T47D cells

**Additional file 7: Figure S1. Growth of ILC and IDC cell lines in 2D and ULA culture. a-b** Relative growth over day 0 of the (**a**) ILC (red) cell lines MM134 (top) and SUM44 (bottom) and (**b**) IDC (blue) cell lines MCF7 (top) and T47D (bottom) in 2D (purple) or ULA (green) culture. Values are from FluoReporter Blue dsDNA assay (n=3). **c** Relative growth over day 0 of the IDC cell line SKBR3 in 2D or ULA culture. Values are from CellTiter-Glo assay (n=6). p-values are from two-way ANOVA comparison of 2D and ULA. * $p \leq 0.05$; *** $p \leq 0.001$; **** $p \leq 0.0001$

**Additional file 7: Figure S2**. **Anoikis resistance of additional ILC and IDC cell lines. a-b** Annexin V and PI FACS staining plots of (**a**) MM330 (top; red) and BCK4 (bottom; red) and (**b**) MM231 (top; blue) and SKBR3 (bottom; blue) cells after 4 days in 2D (left; purple) or ULA (right; green) culture. **c-d** Quantification of the anoikis resistant (Annexin V-/PI-) population in (C) ILC and (D) IDC cell lines. Data is displayed as relative to the 2D condition in each cell line. **e** Immunoblotting for PARP in ILC and IDC cell lines after 2 days in 2D or ULA culture. STAU:

positive control from MM231 cells treated with 1 μM Staurosporine for 5 hours. β-Actin was used as a loading control

**Figure S3. Ki67 levels in ILC and IDC cell lines in 2D and ULA culture as a read-out of cell proliferation. a-b** Representative FACS plots from Ki67 staining of the (**a**) ILC (red) cell lines MM134 (top) and SUM44 (bottom) and (**b**) IDC (blue) cell lines MCF7 (top) and T47D (bottom) after 4 days in 2D (left; purple) or ULA (right; green) culture. Gates were placed based on isotype staining in each cell line in each condition. **c-d** Quantification of the Ki67+ cells based on the gating in (**a-b**) in (**c**) ILC and (**d**) IDC cell lines. Data is displayed as mean percentage +/- standard deviation (n=3). p-values are from t-tests. * $p \leq 0.05$; ** $p \leq 0.01$

**Figure S4. Effects of ROCK inhibition on the viability and morphology of additional ILC and IDC cell lines in 2D and ULA culture. a-b** Dose response curves of the (**a**) ILC (red) cell lines MM330 (left) and BCK4 (right) and (**b**) IDC (blue) cell lines MM231 (left) and SKBR3 (right) treated with the indicated doses of the Y-27632 ROCK inhibitor in 2D (purple) or ULA (green) culture after 4 days. Arrows indicate the dose used for the morphology pictures in (**c-d**). **c** Morphologies of the (**c**) ILC and (**d**) IDC cell lines from c treated with vehicle or 10 μM ($10^{-5}$ M) Y-27632 for 4 days in 2D or ULA culture. Insets show higher magnification images. Scale bar: 100 μm

**Figure S5. Effects of ROCK, p120 and YAP knockdown on the viability of ILC cell lines in 2D and ULA culture. a-b** Immunoblotting for p120, ROCK1 and YAP (**a**) and relative growth over day 0 in 2D or ULA culture (**b**) in the ILC cell lines MM134 (left; top) and SUM44 (right; bottom) transiently transfected with a control, ROCK1, p120 or YAP siRNA. β-Actin was used as a loading control. Graphs show mean +/- standard deviation (n=6). p-values are from ordinary one-way ANOVA with Dunnett's multiple comparison test to siControl in 2D or ULA separately

**Figure S6. Full RPPA heat map**

**Figure S7. Additional proteomic analysis and drug treatments of ILC and IDC cell lines in 2D and ULA culture. a** Western blot analysis of the ILC (red) cell lines MM134 and SUM44 grown in 2D (purple) or ULA (green) culture for 4 days for the indicated pathways proteins. Three biological replicates are displayed for each condition. β-Actin was used as a loading control. **b** Dose response curves of the ILC (red) cell lines MM134 and SUM44 and IDC (blue) cell lines MCF7 and T47D treated with the indicated doses of the p90-RSK inhibitor LJH-685 in 2D (purple; top) or ULA (green; bottom) culture for 4 days

**Figure S8. Transcriptomic profiling of ILC and IDC cell lines in 2D and ULA culture. a** Venn diagrams showing the overlap between the genes downregulated after 24 hours in ULA (green) culture as compared to 2D (purple) in ILC (red) and IDC (blue) cell lines. The list on the right shows the 24 genes commonly downregulated in the two ILC but not the IDC cell lines, mostly made up of small nuclear, nucleolar and spliceosomal RNAs. **b-c** qRT-PCR validation of the ID1 (left) and ID3 (right) knockdown in MM134 (**b**) and SUM44 (**c**) cells 4 days after transient transfection with the indicated siRNAs. Data is displayed as mean +/- error relative to siControl in each condition in each cell line

Figure S1

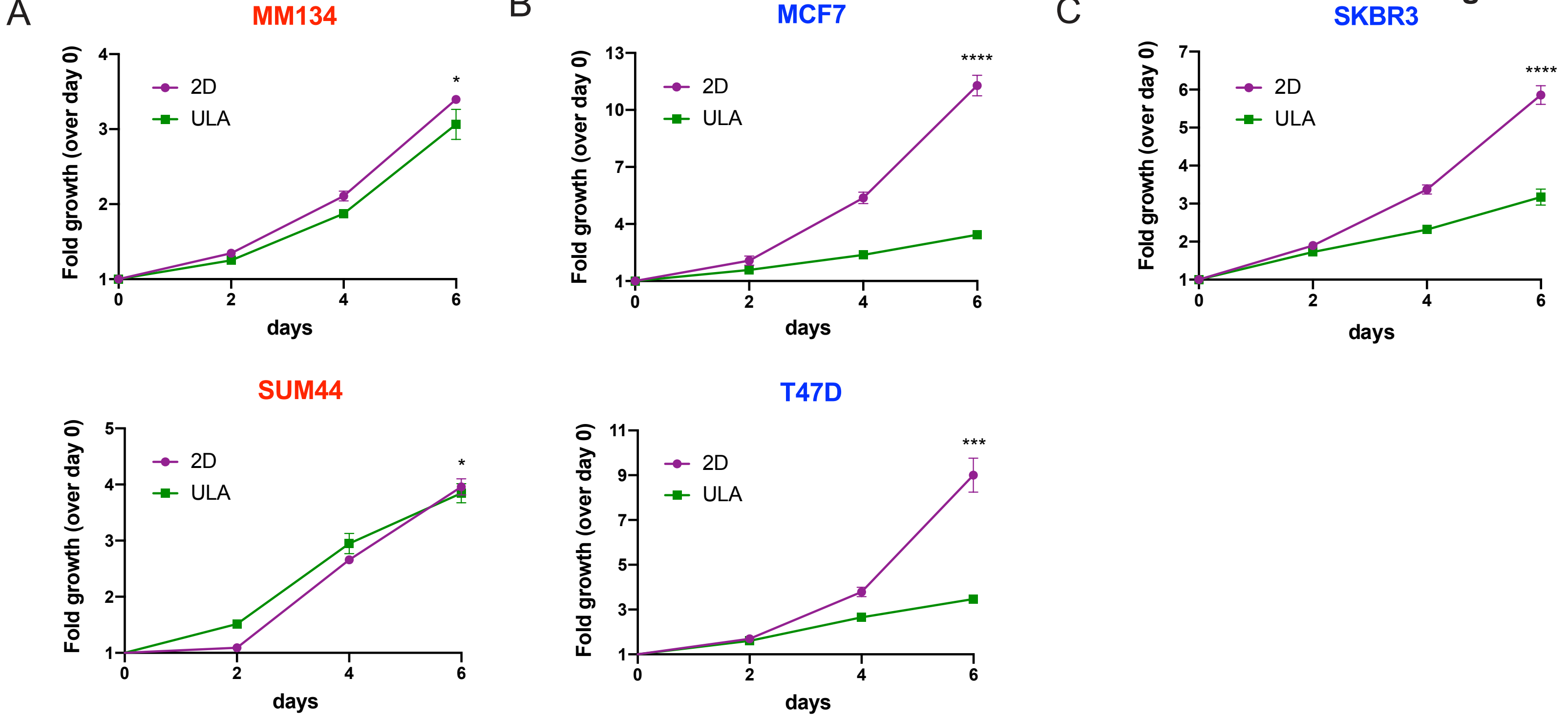


**Figure S1. Growth of ILC and IDC cell lines in 2D and ULA culture. a-b** Relative growth over day 0 of the (**a**) ILC (red) cell lines MM134 (top) and SUM44 (bottom) and (**b**) IDC (blue) cell lines MCF7 (top) and T47D (bottom) in 2D (purple) or ULA (green) culture. Values are from Fluoreporter Blue dsDNA assay (n=3).
**c.** Relative growth over day 0 of the IDC cell line SKBR3 in 2D or ULA culture. Values are from CellTiter-Glo assay (n=6). p-values are from two-way ANOVA comparison of 2D and ULA. * $p \leq 0.05$; *** $p \leq 0.001$; **** $p \leq 0.0001$

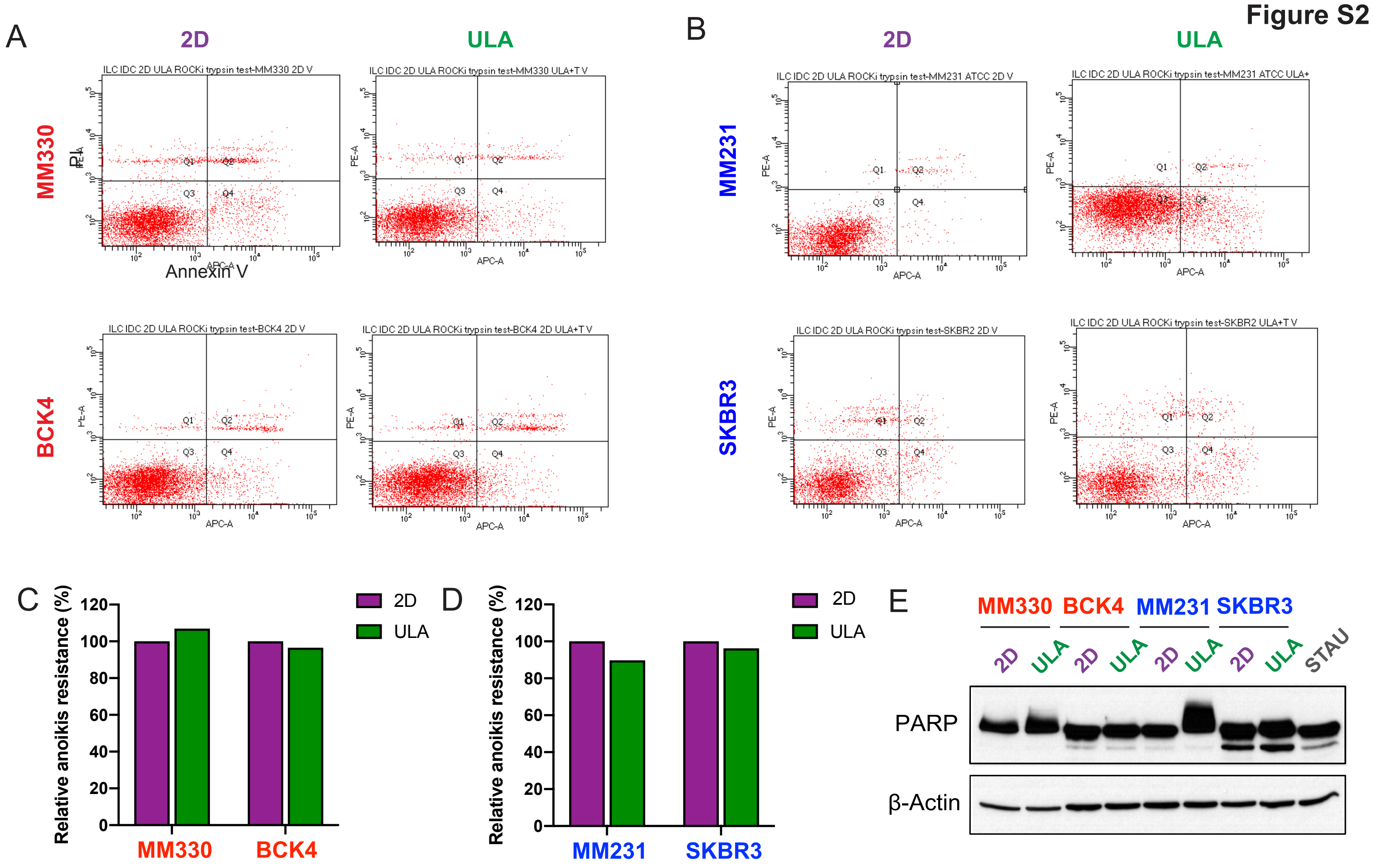


**Figure S2. Anoikis resistance of additional ILC and IDC cell lines. a-b** Annexin V and PI FACS staining plots of (**a**) MM330 (top; red) and BCK4 (bottom; red) and (**b**) MM231 (top; blue) and SKBR3 (bottom; blue) cells after 4 days in 2D (left; purple) or ULA (right; green) culture. **C-D.** Quantification of the anoikis resistant (Annexin V-/PI-) population in (**C**) ILC and (**D**) IDC cell lines. Data is displayed as relative to the 2D condition in each cell line. **E.** Immunoblotting for PARP in ILC and IDC cell lines after 2 days in 2D or ULA culture. STAU: positive control from MM231 cells treated with 1 μM Staurosporine for 5 hours. β-Actin was used as a loading control.

Figure S3

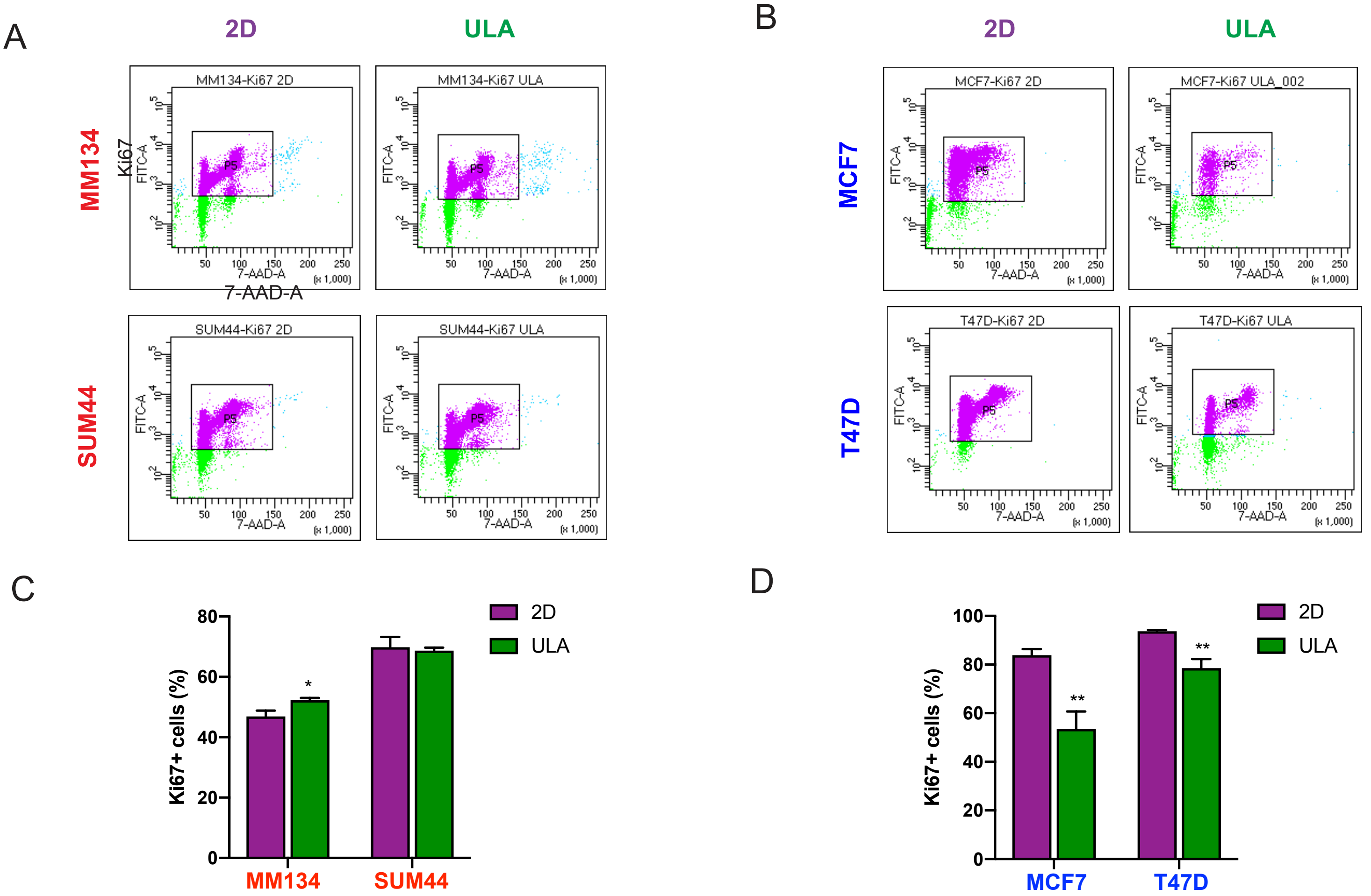


**Figure S3. Ki67 levels in ILC and IDC cell lines in 2D and ULA culture as a read-out of cell proliferation.** **a-b** Representative FACS plots from Ki67 staining of the (**a**) ILC (red) cell lines MM134 (top) and SUM44 (bottom) and (**b**) IDC (blue) cell lines MCF7 (top) and T47D (bottom) after 4 days in 2D (left; purple) or ULA (right; green) culture. Gates were placed based on isotype staining in each cell line in each condition. **c-d** Quantification of the Ki67+ cells based on the gating in (**a-b)** in (**c**) ILC and (**d**) IDC cell lines. Data is displayed as mean percentage +/- standard deviation (n=3). p-values are from t-tests. * $p \leq 0.05$; ** $p \leq 0.01$

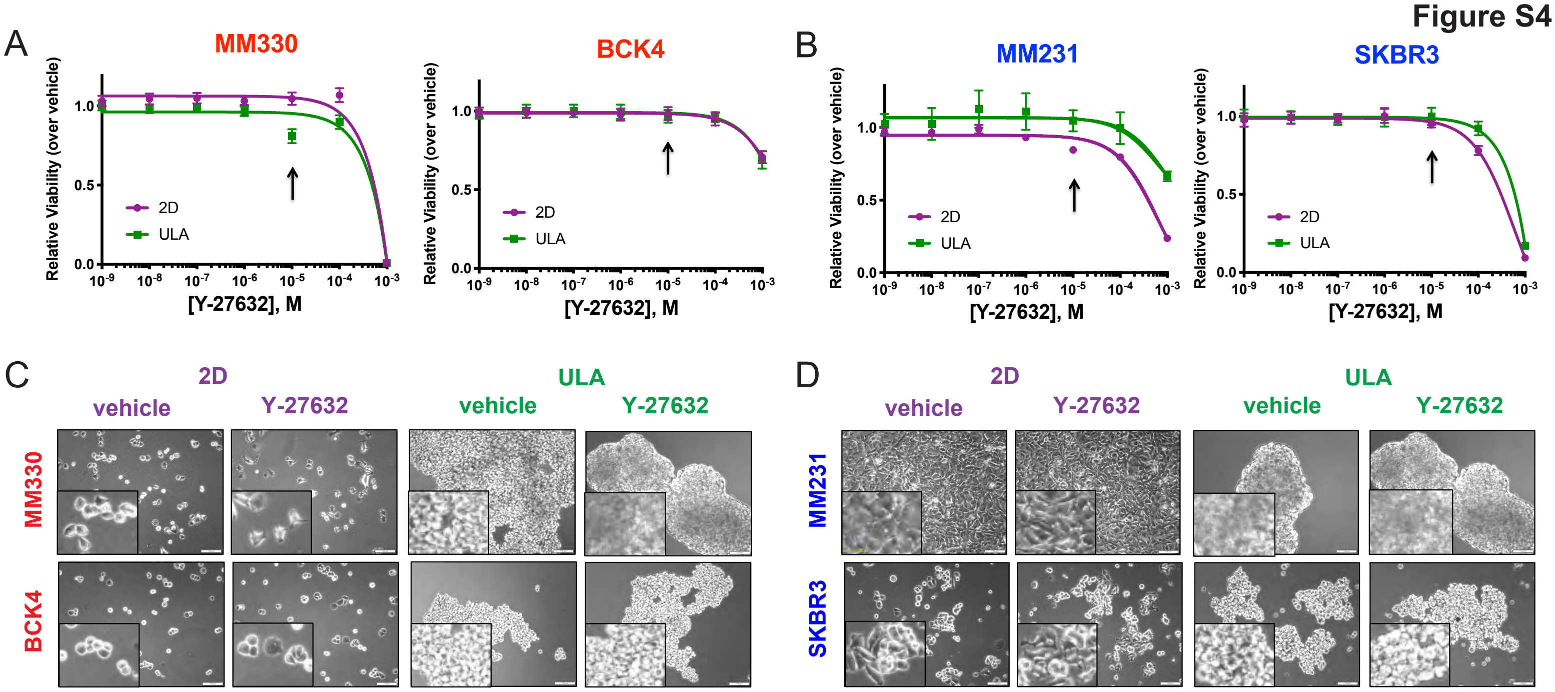


**Figure S4. Effects of ROCK inhibition on the viability and morphology of additonal ILC and IDC cell lines in 2D and ULA culture. a-b** Dose response curves of the (**a**) ILC (red) cell lines MM330 (left) and BCK4 (right) and (**b**) IDC (blue) cell lines MM231 (left) and SKBR3 (right) treated with the indicated doses of the Y-27632 ROCK inhibitor in 2D (purple) or ULA (green) culture after 4 days. Arrows indicate the dose used for the morphology pictures in (**c-d**). **c** Morphologies of the (**c**) ILC and (**d**) IDC cell lines from c treated with vehicle or 10 µM ($10^{-5}$ M) Y-27632 for 4 days in 2D or ULA culture. Insets show higher magnification images.
Scale bar: 100 µm

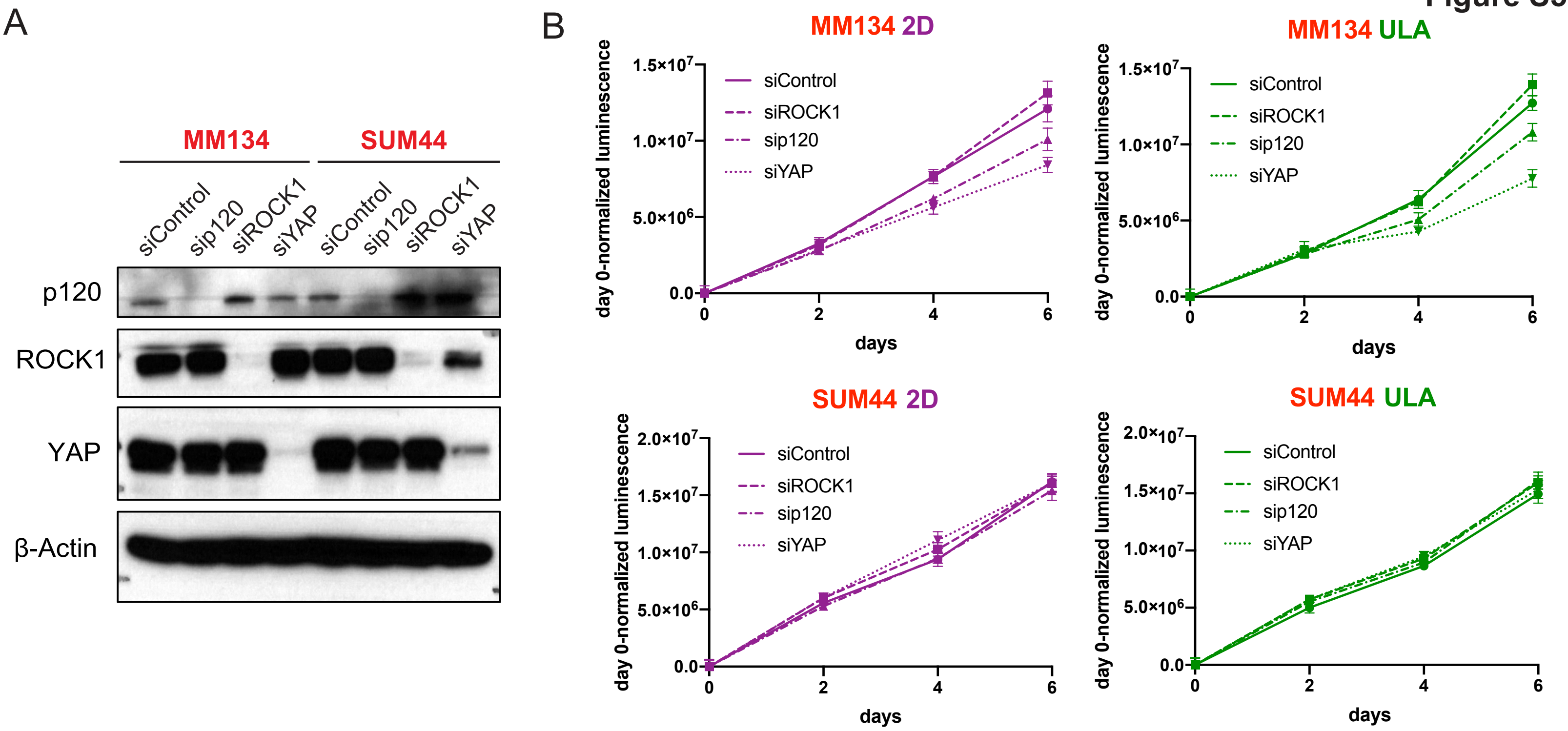


**Figure S5. Effects of ROCK, p120 and YAP knockdown on the viability of ILC cell lines in 2D and ULA culture. a-b** Immunoblotting for p120, ROCK1 and YAP (**a**) and relative growth over day 0 in 2D or ULA culture (**b**) in the ILC cell lines MM134 (left; top) and SUM44 (right; bottom) transiently transfected with a control, ROCK1, p120 or YAP siRNA. β-Actin was used as a loading control. Graphs show mean +/- standard deviation (n=6). p-values are from ordinary one-way ANOVA with Dunnett's multiple comparison test to siControl in 2D or ULA separately

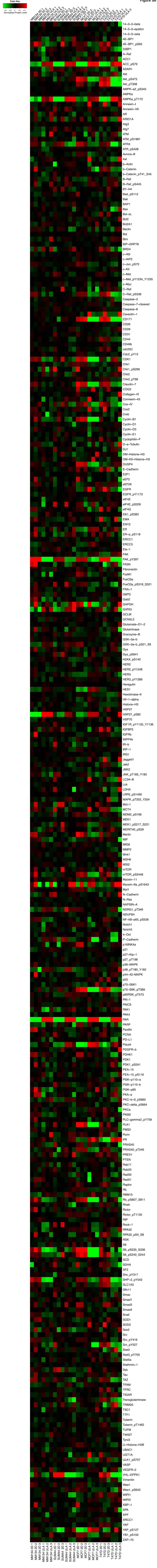

Figure S6. Full RPPA heat map

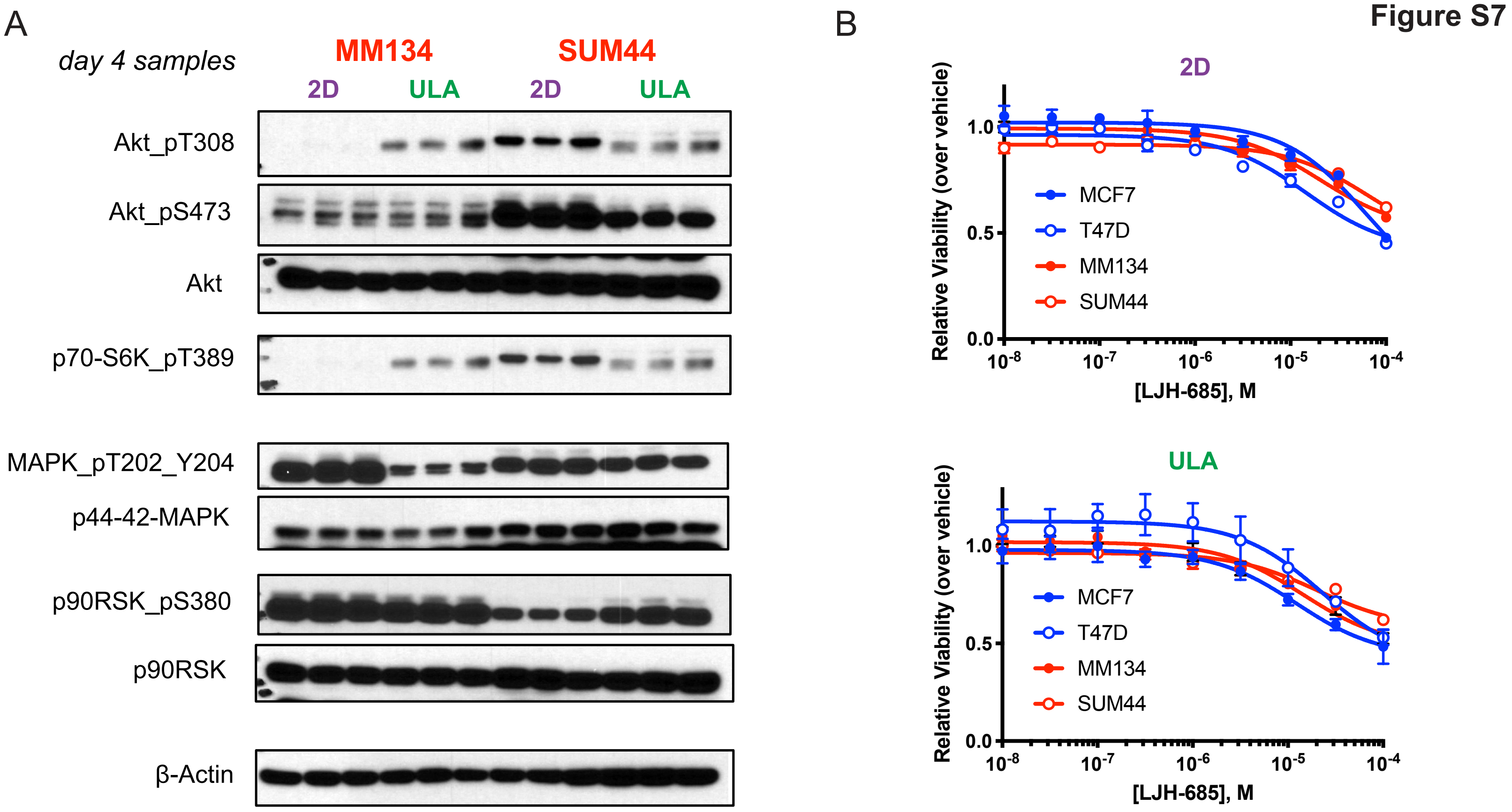


**Figure S7. Additional proteomic analysis and drug treatments of ILC and IDC cell lines in 2D and ULA culture. a** Western blot analysis of the ILC (red) cell lines MM134 and SUM44 grown in 2D (purple) or ULA (green) culture for 4 days for the indicated pathways proteins. Three biological replicates are displayed for each condition. β-Actin was used as a loading control. **b** Dose response curves of the ILC (red) cell lines MM134 and SUM44 and IDC (blue) cell lines MCF7 and T47D treated with the indicated doses of the p90-RSK inhibitor LJH-685 in 2D (purple; top) or ULA (green; bottom) culture for 4 days

Figure S8

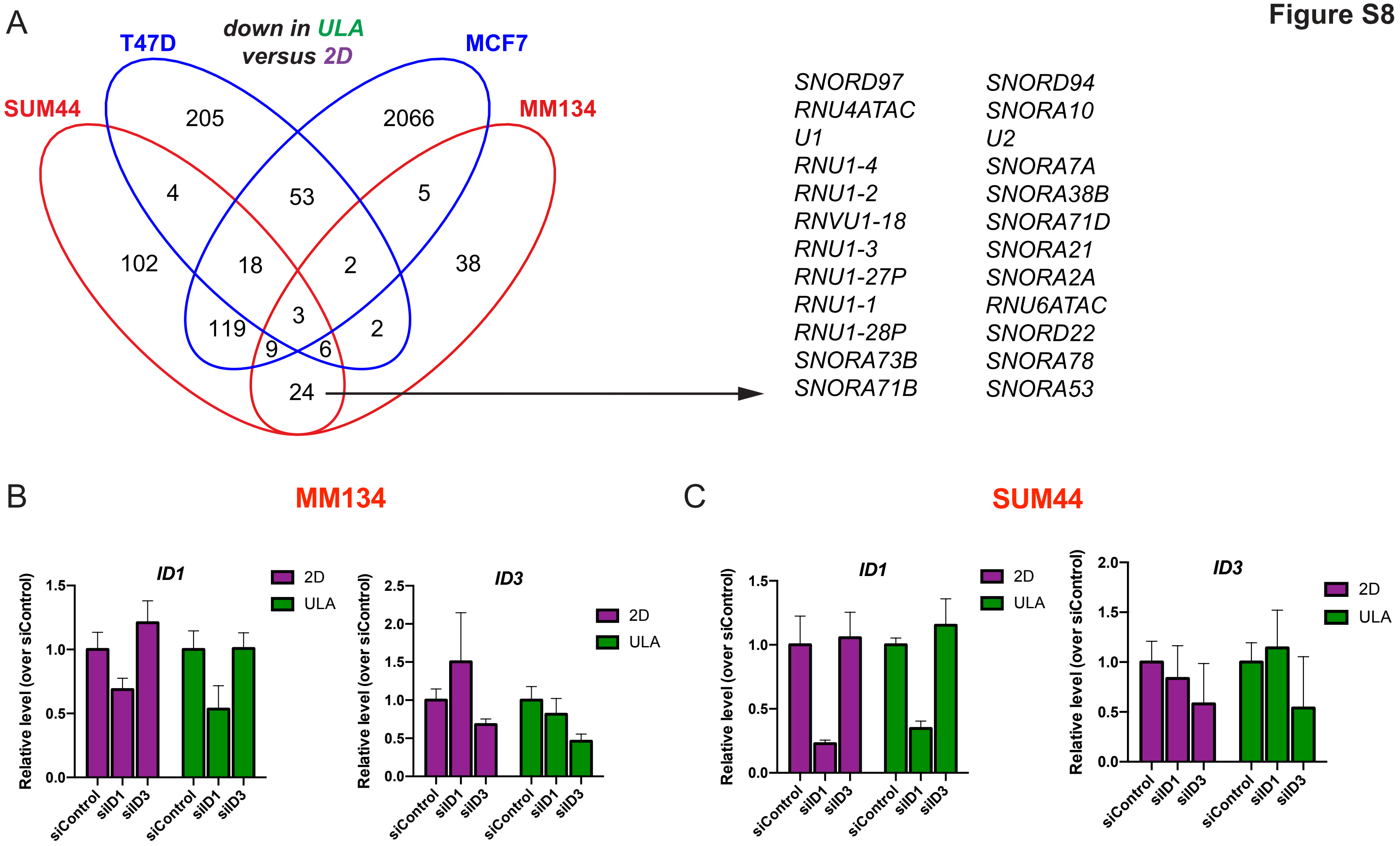


**Figure S8. Transcriptomic profiling of ILC and IDC cell lines in 2D and ULA culture. a** Venn diagrams showing the overlap between the genes downregulated after 24 hours in ULA (green) culture as compared to 2D (purple) in ILC (red) and IDC (blue) cell lines. The list on the right shows the 24 genes commonly downregulated in the two ILC but not the IDC cell lines, mostly made up of small nuclear, nucleolar and spliceosomal RNAs. **b-c** qRT-PCR validation of the *ID1* (left) and *ID3* (right) knockdown in MM134 (**b**) and SUM44 (**c**) cells 4 days after transient transfection with the indicated siRNAs. Data is displayed as mean +/- error relative to siControl in each condition in each cell line
